# An organotypic hippocampo-prefrontal co-culture model for studying long-range connectivity and circuit plasticity

**DOI:** 10.64898/2026.10.20.760475

**Authors:** Phyllis Stöhr, Gayane Grigoryan, Maximilian Lenz, Andreas Vlachos

## Abstract

The hippocampal projection to the medial prefrontal cortex (mPFC) is a key component of brain networks involved in cognition and emotion. Here, we establish and validate an organotypic entorhino-hippocampo-prefrontal co-culture model that enables experimental investigation of long-range circuit connectivity and plasticity under controlled *in vitro* conditions. Anatomical tracing, viral labeling, and ultrastructural analyses demonstrate hippocampal axonal projections and synapse formation within the mPFC. Functional connectivity was supported by hippocampal CA1 stimulation, which evoked postsynaptic responses in 90.6% of recorded layer 2/3 mPFC pyramidal neurons. Intrinsic membrane properties and excitatory synaptic transmission were largely preserved across culture configurations. To demonstrate the experimental utility of the model, we examined the effects of selective alveus lesion and global activity blockade. Alveus lesion produced no detectable changes in the examined synaptic or intrinsic properties of mPFC neurons, whereas tetrodotoxin-mediated activity blockade increased miniature excitatory postsynaptic current frequency and reduced membrane capacitance. Together, these findings establish a versatile experimental platform for investigating long-range circuit connectivity and activity-dependent plasticity, with potential applications in mechanistic studies of neuromodulatory interventions.

## INTRODUCTION

The interaction between the hippocampus and the medial prefrontal cortex (mPFC) constitutes a fundamental organizational principle of higher-order brain function and is critically involved in memory, executive control, and emotional processing (1,2). Disrupted hippocampo-prefrontal communication has been implicated in numerous neuropsychiatric disorders (2). While *in vivo* studies have provided important insights into the behavioral relevance of this circuit, the cellular and synaptic mechanisms underlying long-range hippocampo-prefrontal interactions remain incompletely understood.

Organotypic slice cultures provide a versatile experimental platform for studying neuronal connectivity and synaptic plasticity under controlled conditions (3–5). In particular, entorhino-hippocampal slice cultures preserve the major cortical input to the hippocampus and have been widely used to study synaptic integration, activity-dependent plasticity, and network stability (6–9). However, these preparations lack hippocampal output pathways to distant cortical targets, limiting their utility for investigating interactions between hippocampal and neocortical networks. An experimentally accessible model incorporating both hippocampal and prefrontal compartments would therefore provide opportunities to investigate long-range circuit connectivity and its plasticity under controlled conditions.

Here, we establish and validate an organotypic entorhino-hippocampo-prefrontal (EH-mPFC) co-culture model that extends classical entorhino-hippocampal preparations by incorporating a neocortical target region. Using complementary anatomical, ultrastructural, and electrophysiological approaches, we demonstrate the establishment of structural and functional hippocampo-prefrontal connectivity in vitro. Furthermore, we assess the experimental utility of this preparation by examining neuronal responses to selective hippocampal pathway disruption and global activity deprivation. This model provides a platform for mechanistic investigations of long-range circuit interactions and activity-dependent plasticity, with potential applications in studies of neuromodulatory interventions.

## MATERIALS AND METHODS

### Key Resources

All key resources such as antibodies, reagents and software used in this study, including catalog numbers and if applicable Research Resource Identifiers (RRIDs), can be accessed in the Supplement.

### Ethics statement

Animals were maintained under standard housing conditions (12 h light/dark cycle, food and water *ad libitum*). All experimental procedures were performed in accordance with national and institutional guidelines for animal welfare (X-21/01B, X-18/02C). Every effort was made to reduce the number of animals used.

### Protocol for preparation of organotypic tissue cultures

#### 0. Principles

1. All work must be performed according to the applicable institutional safety regulations, including personal protective equipment (PPE).
2. Animal treatment must be conducted in compliance with the regulations set by the respective regulatory authorities. At all times seek to minimize animal distress.
3. To minimize risk of contamination all work including preparation of media must be performed on a disinfected bench with ventilation switched on.
4. All instruments must be appropriately disinfected (either using 70% ethanol or autoclaved depending on the instrument).
5. Slice cultures were prepared from postnatal day 3–5 C57BL/6J and Thy1-eGFP-M (10) mice

#### 1. Equipment and reagents

- instruments

- one pair of fine-tipped forceps (Figure 1A, (2))
- one pair of angled, blunt forceps (Figure 1A, (6))
- two small spatulas (Figure 1B, (18a & 18b))
- four scalpel holders (Figure 1A (7), Figure 1B (17a & 17b) Figure 1C (19))
- two pairs of fine super-cut scissors (Figure 1A, (5a & 5b))
- one pair of large scissors (Figure 1A, (1))
- one 1 ml pipette (Figure 1A, (9))
- one 2.5 ml pipette (Figure 1A, (10))
- one manipulator for specimen plate (Figure 1C, (23))
- two specimen plates (Figure 1C, (20a & 20b))
- one beaker filled with app. 20 ml water (Figure 1A, (15))
- one beaker filled with app. 20 ml 70% ethanol (Figure 1A, (16))
- usables (sterile)

- one small petri dish per mouse (Figure 1A, (13))
- one big petri dish per mouse (Figure 1A, (14))
- one cotton swab (Figure 1A, (4))
- single-use pipette tips 1 ml (Figure 1A, (9))
- single-use pipette tip 2.5 ml (Figure 1A, (10))
- one PTFE filter per mouse (Figure 1A, (11))
- one 6-well plate per maximum of six mice (Figure 1A, (12))
- three curved scalpel blades (Figure 1A, (7), Figure 1B, (17a & 17b))
- one straight scalpel blade (Figure 1C, (19))
- one vibratome blade (Figure 1C, (22))
- histoacryl glue (Figure 1A, (8))
- devices

- bench with ventilation
- microscope with light and magnification of 10x
- vibratome with calibration check and double-walled buffer tray (Figure 1C, (21))

- incubator with HEPA-filter

#### 2. Preparation of media

- The media for slice culture preparation and incubation should be prepared beforehand.
- Preparation medium (e.g. prepare at least 500ml):

- 100 % (v/v) minimal essential medium (MEM)
- 25 mM HEPES buffer solution
- 0.65 % glucose
- 0.1 mg/mL Streptomycin
- 100 U/mL Penicillin
- 2 mM Glutamax
- adjust pH to 7.38-7.4 (using 1M NaOH und 1M HCl) and keep cool at 4°C
- Incubation medium (at least 1ml per mouse/well):

- 50 % (v/v) MEM
- 25 % (v/v) basal medium Eagle (BME)
- 25 % (v/v) heat-inactivated normal horse serum (NHS)
- 25 mM HEPES buffer solution
- 0.15 % (w/v) Bicarbonate
- 0.65 % (w/v) glucose
- 0.1 mg/ml Streptomycin
- 100 U/ml Penicillin
- 2 mM GlutaMAX
- adjust pH to 7.3 (using 1M NaOH und 1M HCl)
- 6-well-plate for incubation:

- fill each well with 1ml of incubation medium, close lid and put into incubator at least one hour before preparation to allow the temperature of the medium to adjust to the temperature of the incubator

#### 3. Setup of Preparation area and slicing station

- Preparation area:

- turn on the laminar flow of the clean bench
- disinfect preparation area including bench with laminar flow, microscope and vibratome
- add blade (Figure 1B, (22)) to vibratome blade holder and perform calibration
- prepare instruments and containers listed above in preparation area and dissection area as depicted in figures 1 and 3
- keep histoacryl glue (Figure 1A, (8)) on ice/ice pack to keep it cool outside the fridge
- cut away the bottom of a single-use 2.5ml tip (Figure 1A, (10)) with clean scissors to achieve an opening with a diameter of app. 0.5 mm and prepare pipette

- prepare animal body bag
- Slicing station:

- fill double-walled buffer tray (Figure 1C, (21)) with preparation medium and set temperature of the buffer tray to 2°C
- lower the blade holder with the blade into the preparation medium to ensure that the medium and the blade are of the same temperature

#### 4. Preparation steps

*The preparation steps should be carried out as quickly as possible while still working precisely to slow down neuronal decay. The preparation should not take longer than 5 minutes in total*.

- Disinfect the skin of the head using a cotton swab (Figure 1A, (4)) soaked in ethanol.
- Decapitate the animal using the big pair of scissors (Figure 1A, (1)), taking care to retrieve the head in a small petri dish (Figure 1A, (13)).
- Discard the animal body in a designated bag.
- Stabilize the head caudally with spatula (Figure 1B, (18a or b)) and pierce the head from rostral-dorsal with fine-tipped forceps (Figure 1A, (2)) to stabilize the skull

**Figure 1.**
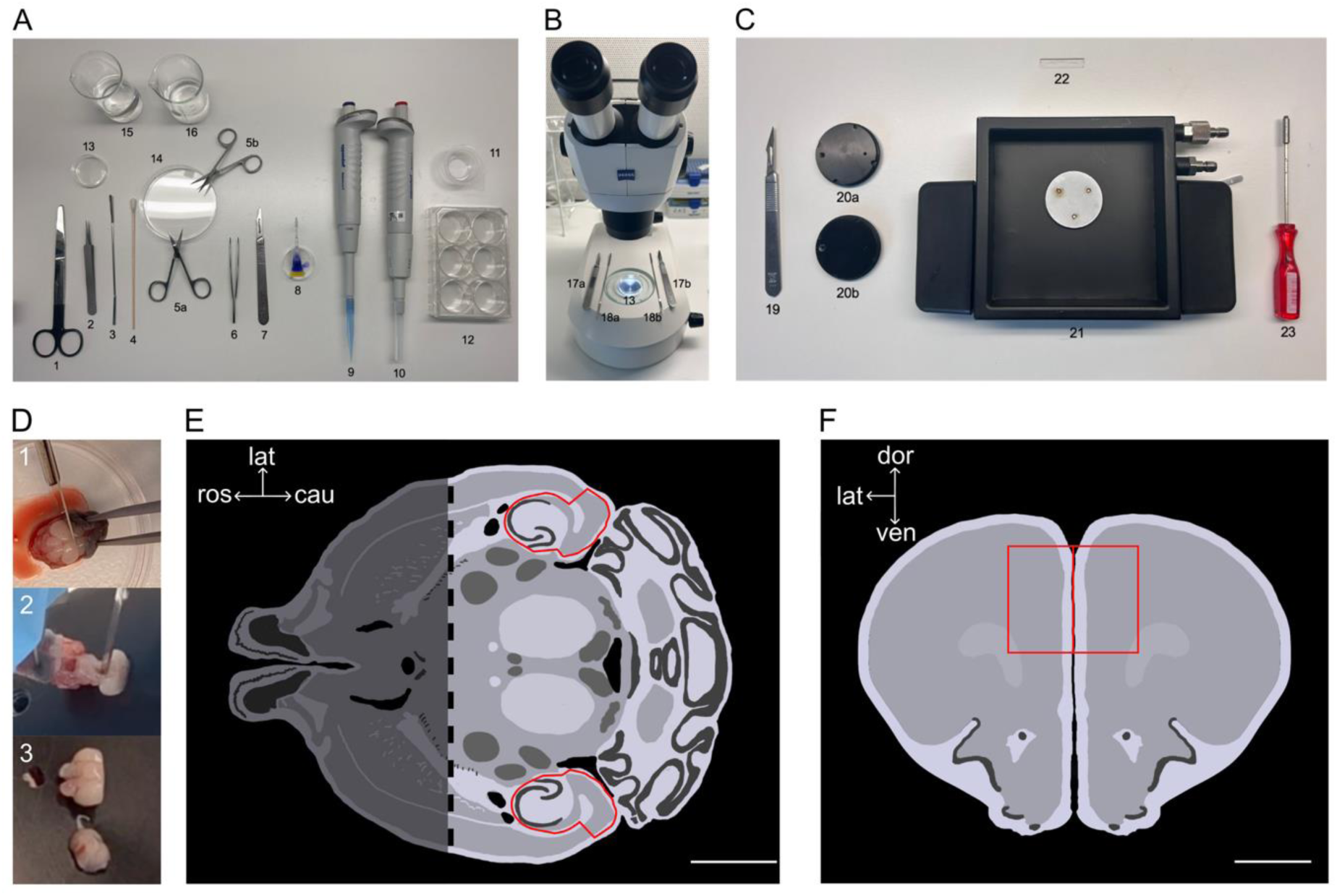
Equipment and critical steps of the preparation process of entorhino-hippocampo-prefrontal (EH-mPFC) co-cultures. **(A)** Instruments and usables necessary for the preparation process labeled 1 through 16 (see *1. Equipment and Reagents*). **(B)** Dissection station with instruments labeled 16 and 17. **(C)** Slicing station with corresponding items labeled 19 through 23 (see *3. Setup of Preparation area and slicing station, Slicing station*). **(D)** Critical steps during the preparation process, for details refer to the Protocol-section: 1 – dissection of rostral mPFC containing brain tissue, 2 – placing caudal brain tissue to the specimen plate, 3 – positioning of both parts of brain tissue on the specimen plate. **(E)** Schematic drawing of a representative horizontal brain slice containing hippocampal formation. The whole slice is shown for context, during the preparation of EH-mPFC co-cultures a cut in frontal plane separates the rostral and the caudal tissue *in situ* (dotted line). Exemplary dissection borders of the entorhino-hippocampal area are indicated with the red line. The orientation of the horizontal slice is provided using directional axes: rostral (ros), caudal (cau) and lateral (lat) directions are provided. Scale bar: 2 mm. **(F)** Schematic drawing of a representative coronal brain slice. Exemplary dissection borders of the mPFC area are indicated with the red line. The orientation of the coronal slice is provided using directional axes: dorsal (dor), ventral (ven) and lateral (lat) directions are provided. Scale bar: 1 mm. Images were adapted from ‘Mouse Brain Atlas: C57BL/6J Horizontal’, Marine Biological Laboratory, https://www.mbl.org/atlas232/atlas232_frame.html. Accessed on 08/28/2026.

*Careful! keep forceps in neutral position, don’t apply closing or opening force in order not to squeeze the tissue)*.

- Keep the forceps with the skull attached to it in your non-dominant hand.
- Using a cotton swab (Figure 1A, (4)), carefully disinfect the skin with 70% ethanol.

*Careful! For the next steps take care to point the instruments upwards in order not to damage the next tissue layer underneath!*

- Incise the skin using fine super-cut scissors (Figure 1A, (5a or b)) in sagittal plane starting caudally and proceeding rostrally until meeting the pointed forceps.
- Make two cuts into the skin angled at 90° to either side of the skull.
- Using angled, blunt forceps (Figure 1A, (6)) peel away the skin and discard it.
- Cut the skull using fine super-cut scissors the same way as the skin previously (first in sagittal plane and afterwards setting two orthogonal cuts).
- Peel away the skull using angled, blunt forceps to expose the brain tissue.
- Use a scalpel with a curved blade (Figure 1A, (7)) to cut through the brain tissue *in situ* in frontal plane approximately 2:3 to achieve a smaller, rostral part containing the mPFC and a bigger, caudal part containing the Hippocampus (Figure 1D, (1)).
- Prepare the specimen plate (Figure 1C, (20a)) by applying a small amount of cold histoacryl glue to the surface.
- Use a small spatula to trace below the caudal tissue part from one side to the other across the base of the skull and tilt the tissue 180° towards the caudal side so the formerly ventral side is flipped to dorsal and *vice versa*. The tissue should remain attached caudally at the cerebellum.
- Still holding the pointed forceps move the head towards the specimen plate.
- Carefully lower the head so the caudal brain tissue touches the histoacryl glue, dorsal side towards the specimen plate.
- Using the spatula carefully pierce through the rostral cerebellum from above to stabilize the tissue. Then pull away the head using the pointed forceps (Figure 1D, (2)). The caudal tissue should now stick to the specimen plate with the ventral surface pointing upwards.
- Add another drop of histoacryl glue app. 1 cm next to the tissue on the specimen plate laterally on one side of your choice.
- Using the spatula carefully trace below the remaining rostral tissue still *in situ* and pull it out of the skull, discard the head afterwards in the animal body bag.
- Guide it onto the other spatula cut side towards the spatula.
- Place it next to the caudal tissue on the histoacryl glue with the cut surface facing downwards and the dorsal part facing in the same direction as the cut surface of the caudal tissue (Figure 1D, (3)).

#### 5. Slicing and dissecting

- Carefully attach the specimen plate inside the buffer tray filled with preparation medium using the manipulator (Figure 1C, (23)). The cut side of the caudal tissue respectively the dorsal side of the rostral tissue should face towards the blade and therefore be cut first.
- Adjust the advance speed and oscillation frequency of the vibratome blade to the desired level, slice the tissue in 300 µm increments.

*Careful! For best results, the cutting process should be carried out very smoothly to avoid tissue damage.*

- Using the 2.5 ml pipette with broadened tip, remove two slices of your choice containing the hippocampus and two slices containing the mPFC and add them to a large petri dish (Figure 1A, (14)) alongside some of the preparation medium.
- Prepare the PTFE-filter (Figure 1A, (11)) by adding 1ml of preparation medium underneath with the 1 ml pipette (Figure 1A, (9)) taking care not to touch the surface.
- Under 10x magnification, using the scalpels with curved blades (Figure 1B, (17a & b)) dissect the entorhino-hippocampal formation from the caudal tissue and the mPFC of the rostral tissue.
- Using the spatulas move the EC-Hipp-tissue-slices to the PTFE-filter.
- Place each mPFC tissue next to one EC-Hipp-tissue with the *alveus* positioned in close proximity to deep mPFC layers (for sectioning strategy and placement of the tissues see Figures 2A,B), thereby approximating the trajectory of hippocampal output pathways(11).

**Figure 2.**
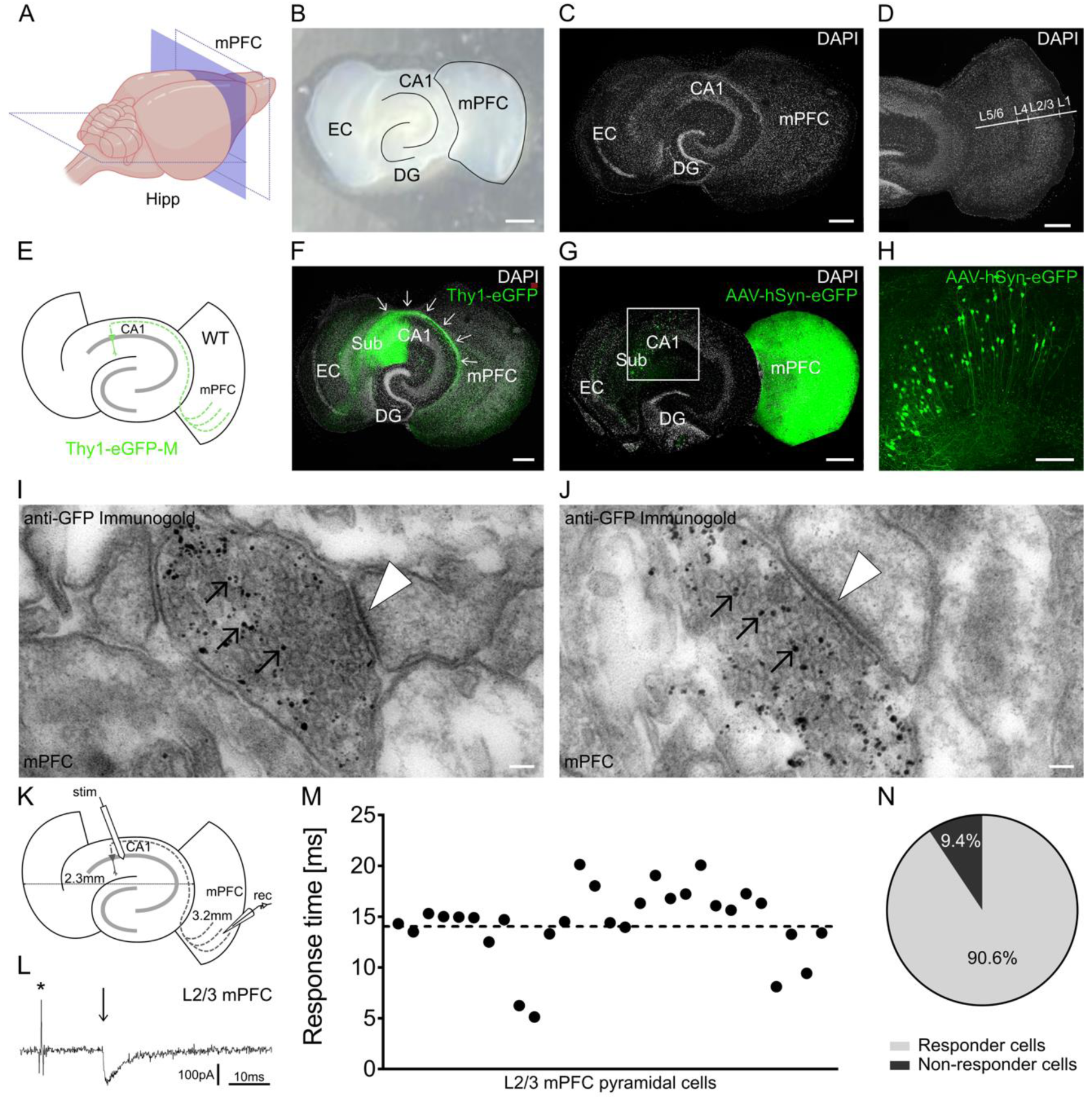
Morphological and functional characterization of organotypic EH-mPFC co-cultures. **(A)** Schematic illustration of the brain dissection strategy. The blue area indicates the cutting plane *in situ*. Dotted lines indicate the planes used for sectioning of the hippocampal formation (Hipp, horizontal) and the medial prefrontal cortex-containing region (mPFC, vertical). **(B)** Representative image of an EH-mPFC co-culture on a membrane insert. Scale bar: 500 µm. EC, entorhinal cortex; DG, dentate gyrus; CA1, hippocampal region cornu ammonis 1. **(C,D)** DAPI staining demonstrating preserved cytoarchitecture of hippocampal subfields and cortical lamination within the mPFC. Scale bars: 300 µm. **(E,F)** Schematic (E) and representative image (F) of an EH-mPFC co-culture generated from Thy1-eGFP-EH and wildtype (WT) mPFC tissue. GFP-positive hippocampal axons (green) extend into the mPFC (*n = 14* cultures). Arrows indicate the trajectory of axonal fibers through the *stratum oriens/alveus* and into prefrontal tissue. Scale bar: 300 µm. **(G,H)** Local AAV2-hSYN-EGFP injection into the mPFC (G) labels hippocampal projection neurons in CA1 and subiculum (H), providing independent evidence for hippocampo-prefrontal connectivity (*n = 4* cultures). Scale bars: 300 µm (G) and 150 µm (H). **(I,J)** Electron micrographs obtained from Thy1-eGFP-M/WT EH-mPFC co-cultures following pre-embedding immunogold labeling for GFP (*n = 6* cultures). GFP-positive presynaptic boutons (indicated with arrows) within the mPFC are opposed to postsynaptic profiles (indicated with arrowheads) and exhibit characteristic synaptic specializations. Scale bar: 50 nm. **(K-N)** Schematic illustration of the recording configuration and distances between stimulation and recording sites (K). Representative postsynaptic response (arrow) recorded from an L2/3 mPFC neuron following extracellular stimulation in hippocampal CA1 (stimulation artifact indicated with asterisk*) (L). Quantification of response latencies (M). Proportion of mPFC neurons exhibiting stimulus-evoked responses following CA1 stimulation (*n = 13* EH-mPFC co-cultures) (N).

#### 6. Incubation

- Remove the prepared 6-well-plate (Figure 1A, (12)) from the incubator and add the PTFE-filter containing EH-mPFC-tissue.
- Return 6-well-plate to the incubator.
- The temperature in the incubator should be set to 35°C, the atmosphere should be humidified with a CO_2_ content of 5%
- Exchange the incubation medium every two days, at least three times per week. Cultures need to be maintained for at least 18 days *in vitro* (DIV) before experimental use.

#### 7. Cleanup and next steps

- Remove the specimen plate from inside the buffer tray and discard the remaining brain tissue, remove histoacryl glue with the scalpel with a straight blade (Figure 1C, (19)).
- After preparing the brain of one animal always clean your dissection instruments in the beakers filled with water (Figure 1A, (15)) and Ethanol (70%) (Figure 1A, (16)) before moving on to the next. Allow Ethanol fully evaporate from the tools before proceeding with next dissection.

#### 8. Culture types

- Three different culture types were generated: entorhino-hippocampo-prefrontal (EH-mPFC) co-cultures as described in the protocol above, entorhino-hippocampal (EH) cultures only containing the entorhino-hippocampal tissue and mPFC cultures only containing the mPFC.

### Whole-cell patch-clamp recordings

Whole-cell patch-clamp recordings were performed as previously described (12–14). Recording electrodes of 3–5 MΩ were used. Patch pipettes were filled with an intracellular solution containing (in mM): 126 K-gluconate, 4 KCl, 4 Mg-ATP, 0.3 Na₂-GTP, 10 phosphocreatine, and 10 HEPES, supplemented with 0.1% (w/v) biocytin. The pH was adjusted to 7.25 with KOH and the osmolarity to 290 mOsm with sucrose. Spontaneous and evoked AMPA receptor-mediated excitatory postsynaptic currents (EPSCs) were recorded from hippocampal CA1 and layer 2/3 (L2/3) mPFC pyramidal neurons at a holding potential of −70 mV. Intrinsic membrane properties were assessed in current-clamp mode using standard current injection protocols. For miniature EPSCs (mEPSCs) recordings, D-APV (10 µM), bicuculline methiodide (10 µM), and TTX (0.5 µM) were added to the recording solution.

mEPSCs were recorded for 2 min. Series resistance was determined before and after each recording. Recordings with a series resistance ≥30 MΩ were excluded from further analysis.

### Extracellular stimulation

Functional hippocampo-prefrontal connectivity was assessed by extracellular stimulation of the CA1 region while recording evoked EPSCs from L2/3 mPFC pyramidal neurons in whole-cell voltage-clamp mode. Monopolar electrical stimuli (1 ms, 3–6 V) were delivered every 10 s through a stimulation electrode positioned at the border between *stratum pyramidale* and *stratum radiatum*. Neurons were classified as exhibiting functional connectivity when reproducible postsynaptic responses were observed in three consecutive stimulation trials. Cells that failed to exhibit responses during four consecutive stimulation trials at the maximal stimulation intensity (6 V) were classified as non-connected. Recordings were terminated once either criterion was met and were excluded if patch-clamp stability was lost before classification. This approach was designed to assess the presence or absence of functional connectivity rather than to quantify stimulus-response relationships.

### Pharmacology

Network activity was suppressed by treatment with TTX (2 µM) for two days prior to electrophysiological recordings as previously described (6,14). Control cultures received vehicle treatment.

### Alveus lesion (AL)

To disrupt major hippocampal output projections, microsurgical transection of the alveus was performed in EH-mPFC co-cultures at 18–20 DIV. Under microscopic control, a scalpel cut was placed at the CA1/CA2 border and extended through the *stratum oriens* to transect the alveus. Cultures were maintained for three days before electrophysiological analysis.

### Post-hoc staining and confocal imaging

Slice cultures were fixed at ≥18 DIV in 4% paraformaldehyde and processed for post-hoc identification of recorded neurons as previously described (12,13,15). Following washing in phosphate buffered saline (PBS), permeabilization and blocking, biocytin-filled neurons were visualized using Alexa Fluor 568-conjugated streptavidin, and DAPI was used for nuclear counterstaining. Cultures were mounted and imaged using a Leica SP8 confocal microscope equipped with 20× and 40× objectives.

### Local transfections

To visualize EH-mPFC projections, cultures were transfected as previously described (9,16) between 3-5 DIV. AAV2-hSYN-EGFP was diluted (1:4) in PBS and approximately 0.1 µl was locally injected into the mPFC. Cultures were returned to the incubator and maintained until at least 18 DIV before analysis.

### Electron microscopy

EH-mPFC co-cultures consisting of EH tissue from Thy1-eGFP-M mice and mPFC tissue from C57BL/6J mice were fixed in 4% PFA and 2% glutaraldehyde (GA) in 0.1 M phosphate buffer (PB) for 1 h, followed by washing in PB. Tissue was sectioned at 50–100 µm and incubated overnight at 4°C with a mouse anti-GFP primary antibody (1:100) in 50 mM Tris-buffered saline containing 2% normal goat serum. For pre-embedding immunogold labeling, sections were incubated overnight with a 1.4 nm Nanogold-conjugated goat anti-mouse secondary antibody (1:100). Following post-fixation in 1% GA, silver enhancement was performed using HQ-Silver. Tissue was subsequently processed for electron microscopy and flat-embedded in Durcupan resin as previously described (12,13).

### Data analysis and statistics

Electrophysiological data were analyzed using pClamp 11 software. Transmission electron microscopy images were analyzed using ImageSP Viewer. Statistical analysis was performed using GraphPad Prism. Data are presented as mean ± SEM. For stimulation delay analyses, measurements were averaged per neuron prior to statistical testing. For all other analyses, individual neurons were considered the experimental unit. Comparisons between two groups were performed using Mann-Whitney-U tests. Statistical significance was set at p < 0.05.

### Data availability

The source data underlying the figures are provided with this manuscript. Additional data supporting the findings of this study are available from the corresponding author upon reasonable request.

### Digital illustrations

ImageJ and Inkscape were used for image processing and figure preparation. Brightness and contrast were adjusted uniformly.

## RESULTS

### Establishment of an organotypic EH-mPFC co-culture

To establish an organotypic *in vitro* model incorporating both hippocampal input and output connectivity with a neocortical target region, we combined organotypic entorhino-hippocampal slices with mPFC slices. mPFC-containing slices were positioned adjacent to the hippocampal formation at the level of the *alveus* (Figures 2A-C), thereby extending the preparation beyond intrinsic and recurrent entorhino-hippocampal connectivity. Following at least 18 DIV, both hippocampal and prefrontal regions remained readily identifiable within EH-mPFC co-cultures. DAPI staining revealed preserved hippocampal cytoarchitecture and cortical lamination within the mPFC, including the expected low cell density of L1 and a discernible boundary to the underlying L2/3 (Figure 2D).

### Anatomical verification of hippocampo-prefrontal projections

To visualize hippocampal projections to the mPFC within EH-mPFC co-cultures, EH slices derived from Thy1-eGFP-M mice were combined with mPFC slices obtained from C57BL/6J mice (Figure 2E). In these cultures, eGFP expression is observed in pyramidal neurons of the CA1 region and subiculum. This configuration enabled selective visualization of hippocampal output projections extending into the mPFC. GFP-positive axons extended from the hippocampal formation into the mPFC in all examined cultures (*n = 14*), where labeled fibers and putative presynaptic boutons were observed throughout the prefrontal neuropil (Figure 2F).

As a complementary approach, AAV2-hSYN-EGFP was locally injected into the mPFC of EH-mPFC co-cultures prepared from C57BL/6J mice (Figure 2G). Following viral expression, GFP-positive neurons were detected in hippocampal regions, including CA1 and the subiculum, in all examined cultures (*n = 4*; Figure 2H).

To determine whether these projections established synaptic contacts, we performed pre-embedding immunogold labeling for GFP followed by electron microscopy in EH-mPFC co-cultures generated from Thy1-eGFP-M and wildtype tissue as described above. Ultrastructural analyses identified GFP-positive presynaptic boutons within the mPFC that were opposed to postsynaptic structures and exhibited characteristic synaptic specializations (Figures 2I,J), consistent with the formation of synaptic contacts between hippocampal projections and prefrontal neurons, in all examined cultures (*n = 6*). Together, these complementary anatomical approaches demonstrate that hippocampal axons extend into the mPFC and establish morphologically identifiable synaptic contacts within the prefrontal tissue.

### Functional verification of hippocampo-prefrontal connectivity

To determine whether established hippocampal projections are providing functional connectivity with the mPFC, whole-cell patch-clamp recordings were obtained from L2/3 mPFC neurons while extracellular stimulation was delivered to the CA1 region (Figure 2K).

Electrical stimulation of CA1 reliably evoked postsynaptic responses in 90.6% of recorded mPFC neurons (29 of 32 cells; Figures 2L–N). Across 13 independent EH-mPFC co-cultures, only three neurons failed to exhibit detectable stimulus-evoked responses. Evoked responses occurred with reproducible latencies relative to the stimulation artifact (Figure 2M, N), consistent with functional communication between the hippocampal and prefrontal compartments. Together with the anatomical and ultrastructural evidence, these findings support the establishment of robust functional hippocampo-prefrontal connectivity in EH-mPFC co-cultures.

### Baseline electrophysiological properties of CA1 and L2/3 mPFC pyramidal neurons

To assess whether co-culturing affects the baseline electrophysiological properties of hippocampal and prefrontal neurons, we compared intrinsic membrane characteristics and excitatory synaptic transmission between conventional organotypic cultures and EH-mPFC co-cultures. Electrophysiological properties of CA1 pyramidal neurons in EH cultures were compared with those recorded in EH-mPFC co-cultures, while L2/3 mPFC pyramidal neurons from isolated mPFC cultures were compared with those from EH-mPFC co-cultures (Figures 3A-F). All recordings were obtained from age-matched cultures at identical DIV and analyzed in parallel experimental sessions.

**Figure 3.**
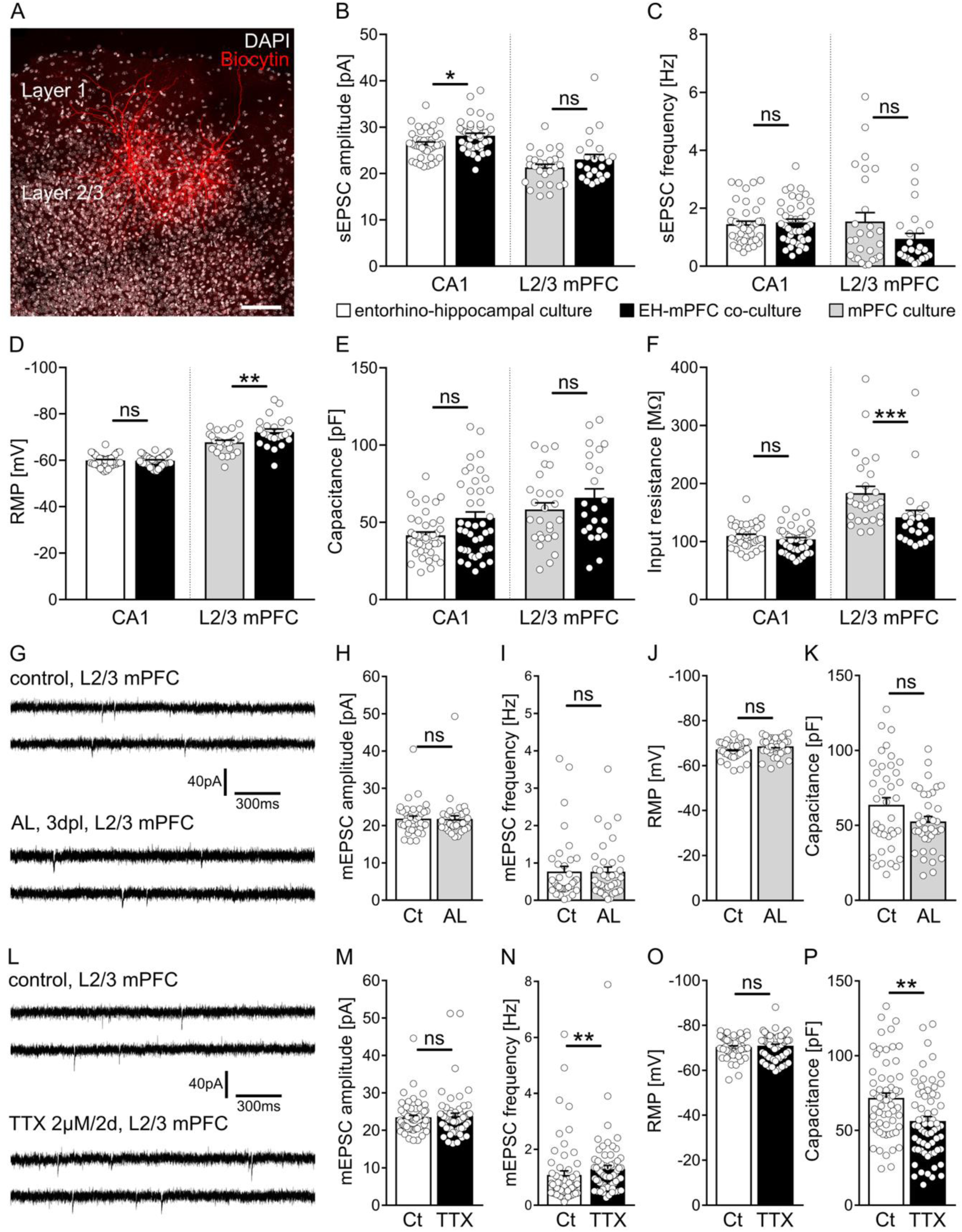
Electrophysiological characterization of EH-mPFC co-cultures under baseline conditions and following selective or global activity deprivation. **(A)** Representative biocytin-filled L2/3 mPFC pyramidal neurons (red) counterstained with DAPI (white). Scale bar: 100 µm. **(B-F)** Group data of AMPA receptor-mediated spontaneous excitatory postsynaptic currents (sEPSCs) and intrinsic membrane properties recorded from hippocampal CA1 pyramidal neurons in entorhino-hippocampal cultures (*n = 42* cells, *13* cultures) and EH-mPFC co-cultures (*n = 41* cells, *10* cultures), as well as from L2/3 mPFC pyramidal neurons in isolated mPFC cultures (*n = 27* cells, *10* cultures) and EH-mPFC co-cultures (*n = 23* cells, *8* cultures). Quantification of sEPSC amplitude (B), sEPSC frequency (C), resting membrane potential (RMP; D), membrane capacitance (E), and input resistance (F). **(G-K)** Examination of synaptic and intrinsic properties of L2/3 mPFC cells following *alveus* lesion (AL). Representative mEPSC recordings obtained from L2/3 mPFC pyramidal neurons under control conditions (Ct; *n = 39* cells, *9* cultures) and three days following AL (3 days post lesion (dpl); *n = 37* cells, *10* cultures) (G). Group data of miniature EPSCs (mEPSCs) and intrinsic membrane properties. Quantification of mEPSC amplitude (H) frequency (I), RMP (J), and membrane capacitance (K). **(L-P)** Effects of global activity deprivation induced by tetrodotoxin (TTX; 2 µM, two days). Representative mEPSC recordings obtained from L2/3 mPFC pyramidal neurons under control conditions (Ct; *n = 60* cells, *13* cultures) and following TTX treatment (*n = 65* cells, *13* cultures) (L). Quantification of mEPSC amplitude (M), frequency (N), RMP (O), and membrane capacitance (P); ns – non significant; * - p < 0.05; ** - p < 0.01; *** - p < 0.001.

Analysis of spontaneous excitatory synaptic transmission revealed largely preserved baseline properties across different culture configurations. In CA1 pyramidal neurons, sEPSC amplitude was modestly increased in EH-mPFC co-cultures compared with EH cultures, whereas sEPSC frequency remained unchanged (Figures 3B,C). In L2/3 mPFC pyramidal neurons, neither sEPSC amplitude nor frequency differed significantly between isolated mPFC cultures and EH-mPFC co-cultures (Figures 3B,C).

Analysis of intrinsic membrane properties demonstrated similarly stable baseline electrophysiological characteristics. In CA1 pyramidal neurons, resting membrane potential (RMP), input resistance (R_in_), and membrane capacitance did not differ between culture configurations (Figures 3D-F). In L2/3 mPFC pyramidal neurons, neurons in EH-mPFC co-cultures exhibited a more hyperpolarized RMP and lower R_in_ compared with isolated mPFC cultures, whereas membrane capacitance remained unchanged (Figures 3D-F).

Together, these findings indicate that the majority of the examined electrophysiological properties are preserved in EH-mPFC co-cultures, although specific differences in synaptic and intrinsic membrane characteristics were observed.

### Effects of selective interruption of hippocampal output on prefrontal neuronal properties

To examine the effects of targeted disruption of hippocampal output pathways, we performed *alveus* lesions (AL) in EH-mPFC co-cultures (Figures 3G–K). Electrophysiological recordings were obtained from L2/3 mPFC pyramidal neurons three days after AL and compared with recordings from non-lesioned cultures. Analysis of mEPSCs revealed no significant effects of AL on excitatory synaptic transmission. Thus, neither mEPSC amplitude nor mEPSC frequency differed significantly between groups (Figures 3H,I).

Intrinsic membrane properties were similarly preserved. RMP and membrane capacitance did not differ significantly in neurons from AL and control EH-mPFC co-cultures (Figures 3J,K).

These findings indicate that *alveus* lesion does not produce detectable changes in the examined excitatory synaptic and intrinsic membrane properties of L2/3 mPFC pyramidal neurons within three days after lesion.

### Effects of global activity deprivation on synaptic and intrinsic properties of mPFC neurons

To determine whether global suppression of neuronal activity induces adaptive changes within EH-mPFC co-cultures, cultures were treated with TTX (2 µM) for two days to block action potential generation and propagation. Electrophysiological recordings were subsequently obtained from L2/3 mPFC pyramidal neurons and compared with those from vehicle-treated control cultures (Figures 3L-P).

TTX treatment significantly altered excitatory synaptic transmission in mPFC neurons. In particular, mEPSC frequency increased following activity blockade, whereas mEPSC amplitude remained unchanged (Figures 3M,N).

Global activity blockade also affected intrinsic membrane properties. Membrane capacitance was significantly reduced in TTX-treated neurons compared with controls, whereas RMP remained unchanged (Figures 3O,P).

Together, these findings demonstrate that global activity deprivation induces measurable changes in excitatory synaptic transmission and intrinsic membrane properties of superficial mPFC neurons within EH-mPFC co-cultures. These results further establish the suitability of the preparation for investigating activity-dependent neuronal adaptations under experimentally controlled conditions.

## DISCUSSION

In this study, we established and functionally validated an organotypic EH-mPFC co-culture model that enables direct investigation of long-range hippocampo-prefrontal connectivity at cellular and synaptic resolution. By combining classical EH slice cultures with mPFC tissue, we generated a preparation in which anatomically defined hippocampal projections establish functional connectivity with prefrontal neurons. This approach extends established organotypic culture models beyond recurrent hippocampal circuitry (4,5,17) and provides experimental access to long-range hippocampo-prefrontal interactions under highly controlled conditions.

Multiple independent lines of evidence support the establishment of functional hippocampo-prefrontal connectivity in EH-mPFC co-cultures. Anatomical tracing demonstrated directed hippocampal projections into the mPFC, ultrastructural analyses confirmed the formation of synaptic contacts, and electrophysiological recordings revealed reliable postsynaptic responses in superficial mPFC neurons following hippocampal CA1 stimulation. Together, these findings support the establishment of structurally and functionally integrated hippocampo-prefrontal connections *in vitro*. Importantly, baseline intrinsic and excitatory synaptic properties remained largely stable across culture configurations, indicating that incorporation of long-range hippocampal projections does not fundamentally alter neuronal physiology within the preparation.

A major advantage of this model is that it permits selective manipulation of long-range connectivity while maintaining experimental accessibility at the level of identified neurons and synapses. As a proof-of-principle application, we examined the consequences of selectively interrupting hippocampal output by *alveus* lesion. Despite robust anatomical and functional connectivity, selective disruption of hippocampal efferents did not induce major alterations in excitatory synaptic transmission or intrinsic membrane properties of superficial mPFC pyramidal neurons within 3 days after the lesion. This finding contrasts with classical denervation paradigms in the hippocampal formation, in which loss of major afferent pathways can trigger pronounced homeostatic adaptations (6,8,13,18,19). The present observations indicate that *alveus* lesion does not elicit detectable changes in the examined electrophysiological properties of superficial mPFC neurons within three days after lesion. However, different forms of synaptic and intrinsic adaptation may exhibit distinct temporal dynamics and region-specific expression, and alterations occurring outside the present observation period or in other brain regions cannot be excluded.

In contrast, global suppression of neuronal activity by TTX induced measurable alterations in both synaptic and intrinsic neuronal properties. Increased mEPSC frequency and reduced membrane capacitance demonstrate that the preparation is responsive to experimentally induced activity deprivation. Although these changes may reflect activity-dependent adaptations, their underlying mechanisms remain to be determined. Together, these findings support the suitability of EH-mPFC co-cultures for studying neuronal responses to defined experimental perturbations (20–22). These findings demonstrate that the preparation is sensitive to network-wide perturbations and therefore suitable for studying activity-dependent plasticity mechanisms across interconnected brain regions.

Beyond these proof-of-principle experiments, EH-mPFC co-cultures provide a platform for investigating how neuronal activity is integrated, propagated, and transformed across anatomically defined long-range circuits. Unlike conventional organotypic hippocampal cultures, the present model incorporates allocortical and neocortical compartments within the same experimentally accessible network. This feature enables direct investigation of hippocampo-prefrontal interactions at cellular and synaptic resolution while preserving the experimental accessibility and long-term stability of organotypic preparations. In addition, the preparation permits selective manipulation of individual circuit components and simultaneous assessment of their impact on connected network regions.

The modular architecture of the preparation further enables the assembly of hybrid circuits from genetically distinct donor animals (c.f., Figure 2E). Such combinations could provide experimental opportunities that are difficult to achieve *in vivo*, including the selective manipulation of individual network nodes while maintaining functional interactions with genetically unmodified partner regions.

The preparation is also highly compatible with emerging circuit-level interrogation technologies. For example, patterned holographic stimulation could be used to selectively activate defined neuronal populations within the entorhinal cortex or hippocampus and thereby probe the propagation of activity across the interconnected network (23). Conversely, modern optical recording approaches, including genetically encoded voltage and calcium indicators, may enable simultaneous monitoring of network dynamics in the respective regions (24,25). Such combinations could facilitate mechanistic studies of activity flow, information transfer, and plasticity across long-range hippocampo-prefrontal circuits under experimentally controlled conditions.

Importantly, the incorporation of neocortical tissue also creates new opportunities for studying the cellular effects of non-invasive neuromodulation. Previous investigations of repetitive magnetic stimulation in organotypic cultures have largely focused on hippocampal circuitry because of the experimental accessibility of archicortical structures (7,9,26). In contrast, most clinical and experimental applications of transcranial magnetic stimulation target neocortical regions (27). The presence of interconnected hippocampal and neocortical compartments may therefore enable the distinction between local and network-mediated effects of neuromodulatory interventions. By incorporating entorhinal, hippocampal, and prefrontal compartments into a single interconnected network, the present preparation enables investigation of stimulation-induced responses across distinct cortical regions under identical experimental conditions. This feature may facilitate mechanistic studies of regional susceptibility, plasticity induction, network interactions, and state-dependent effects of neuromodulatory interventions.

Several limitations should be considered. The preparation does not recapitulate the full complexity of the hippocampo-prefrontal network and lacks subcortical afferents, long-range neuromodulatory inputs, and physiological behavioral states. Moreover, hippocampo-prefrontal connections develop within a reorganized *in vitro* environment and may not fully reproduce the anatomical specificity and functional organization of the corresponding projections *in vivo*. Although anatomical, ultrastructural, and electrophysiological evidence supports the establishment of functional connectivity, the precise synaptic organization and contribution of individual hippocampo-prefrontal projections remain to be determined. Nevertheless, the robust anatomical and functional connectivity observed in the present study supports its utility for mechanistic investigations of long-range interactions under controlled experimental conditions.

Taken together, the organotypic hippocampo-prefrontal co-culture model introduced here provides an experimentally accessible platform for investigating long-range circuit connectivity and activity-dependent plasticity. By combining anatomically interconnected hippocampal and neocortical compartments with electrophysiological accessibility and compatibility with modern circuit interrogation approaches, this preparation offers new opportunities to examine the cellular and synaptic mechanisms underlying interregional neuronal communication. Its modular architecture may further facilitate mechanistic studies of neuromodulatory interventions and their effects on distributed neuronal networks.

## Supporting information

Key Resources Table

## ACKNOWLEDGEMENTS

We thank Mrs. Emina Deumic for excellent technical support and Mrs. Sigrun Nestel for support in electron microscopy.

